# Coordinated regulation of the metaboproteome by Hsp90 chaperones controls metabolic plasticity

**DOI:** 10.1101/2025.11.21.689798

**Authors:** R Felipe Perez, Gianna L Mochi, Julia K Burkacki, Victoria Zoccoli-Rodriguez, Landon D Arcadi, Sarah J Backe, Joel R Wilmore, Mark R Woodford

**Author notes:** These authors contributed equally to this work (R Felipe Perez, Gianna L Mochi).

## Abstract

Heat shock protein 90 (Hsp90) chaperones participate in the stabilization and activation of hundreds of proteins, thereby acting as signaling hubs. A mitochondrial subpopulation of Hsp90 has been previously described; however, little is known about its role in metabolism. Here, we showed that loss of individual Hsp90 isoforms differentially affects oxygen consumption and metabolic flexibility. Proteomic and metabolomic evaluation demonstrated that Hsp90 regulates the mitochondrial metabolic network, including respiration, fatty acid oxidation, and redox homeostasis. Loss of the mitochondrial chaperone TRAP1 induced compensatory binding of Hsp90s to TRAP1-dependent proteins, indicating a mechanism for the role of Hsp90 chaperones in metabolic reprogramming. When considered with previous findings, a temporal pattern of regulation emerges whereby Hsp90s control the transcription, translation, import, and assembly of mitochondrial protein complexes. Our findings expand the scope of Hsp90-regulated processes and potentially inform the effects of isoform-specific Hsp90 inhibitors on metabolic reprogramming in cancer and other diseases.

## INTRODUCTION

Metabolism is governed by the activity, abundance, and localization of enzymes whose functions are intimately linked to their structural integrity. Metabolic homeostasis depends not only on gene expression and allosteric regulation but also on the correct folding, maturation, and turnover of the metabolic proteome^1,2^. Recent studies have highlighted the responsiveness of metabolism to proteostatic stress, including the activation of adaptive pathways such as the unfolded protein response (UPR^3,4^), the integrated stress response (ISR^5,6^), and autophagy^7^. However, the extent to which molecular chaperones control metabolic protein function remains undetermined.

Heat shock protein 90 (Hsp90) is a family of molecular chaperones that use the energy of ATP hydrolysis to regulate the stabilization and activation of their substrate proteins, or ‘clients^8,9^.’ The constitutively expressed Hsp90β and stress-inducible Hsp90α isoforms regulate the function of hundreds of client proteins, thereby acting as hubs of cellular signaling^10^. Hsp90α and Hsp90β collaborate with co-chaperone proteins that deliver substrates, regulate ATP hydrolysis, and direct Hsp90’s structural rearrangements to fine-tune client function^11^. This regulatory framework comprises the ‘chaperone cycle^8,9^’. By regulating client function, Hsp90 has been shown to directly control many cellular processes including mitogen-activated protein kinase (MAPK) signaling^12^, autophagy^13^, and the cell cycle^14,15^.

Hsp90α and Hsp90β are found abundantly in the cytosol, and have been observed to traffic mitochondrial precursor proteins to the translocase of the outer mitochondrial membrane (TOM) complexes for mitochondrial import. The mitochondria-specific Hsp90 isoform, TNF Receptor-Associated Protein 1 (TRAP1), contains an N-terminal mitochondrial targeting signal and has a role in regulating the function of Complexes II and III of the electron transport chain (ETC)^16^; however, the broader impact of TRAP1 activity is poorly understood^17,18^. Previous works in mammalian cells have observed an intramitochondrial population of Hsp90^16,19^ that impacts respiration^20^. Additionally, a role has been attributed to Hsp90 in regulating the mitochondrial permeability transition^21,22^, a precipitating event in certain cell death pathways^23^. Despite this, the interplay between Hsp90α, Hsp90β and TRAP1 in the regulation of mitochondrial metabolism has not been investigated.

In this study, we set out to systematically investigate the impact of individual Hsp90 chaperone knock-outs on the cellular metabolic network. We employed both pharmacological and genetic strategies to observe the effects of chaperone function loss on the metabolome and ‘metaboproteome,’ or the protein network that supports metabolism in the cytosol and mitochondria. By mapping metabolic flux changes in response to specific chaperone perturbations, we find that loss of Hsp90α or Hsp90β reprograms the mitochondrial proteome, resulting in dysregulation of the metabolite landscape spanning both intra- and extra-mitochondrial metabolic pathways.

## RESULTS

### Hsp90s control mitochondrial structure and function

To understand the function of Hsp90 in the mitochondria, we first examined the impact of specific Hsp90α and Hsp90β knockout (KO) on mitochondrial network organization. In the presence of MitoTracker Green, we observed that while mitochondria from wild-type HEK293 cells were small and round, mitochondria from Hsp90α and Hsp90β KO cells were more filamentous and tubular (**Figure 1A**). Further, tetramethylrhodamine, methyl ester (TMRM) uptake indicated that mitochondrial membrane potential was decreased in Hsp90 KO cells (**Figure S1A**). We therefore asked whether the loss of membrane potential could be explained by changes in mitochondrial ultrastructure. Interestingly, we found that mitochondria from Hsp90α KO cells were morphologically distinct from those in WT and Hsp90β KO cells, exhibiting increased mitochondrial number and cristae score (a measure of cristae abundance, definition and organization^24^) (**Figure 1B-C, S1B-D**). Interestingly, mitochondria from Hsp90β KO cells were more similar to WT counterparts, both in gross morphology and cristae score (**Figure 1B-C, S1B-D**).

**Figure 1).**
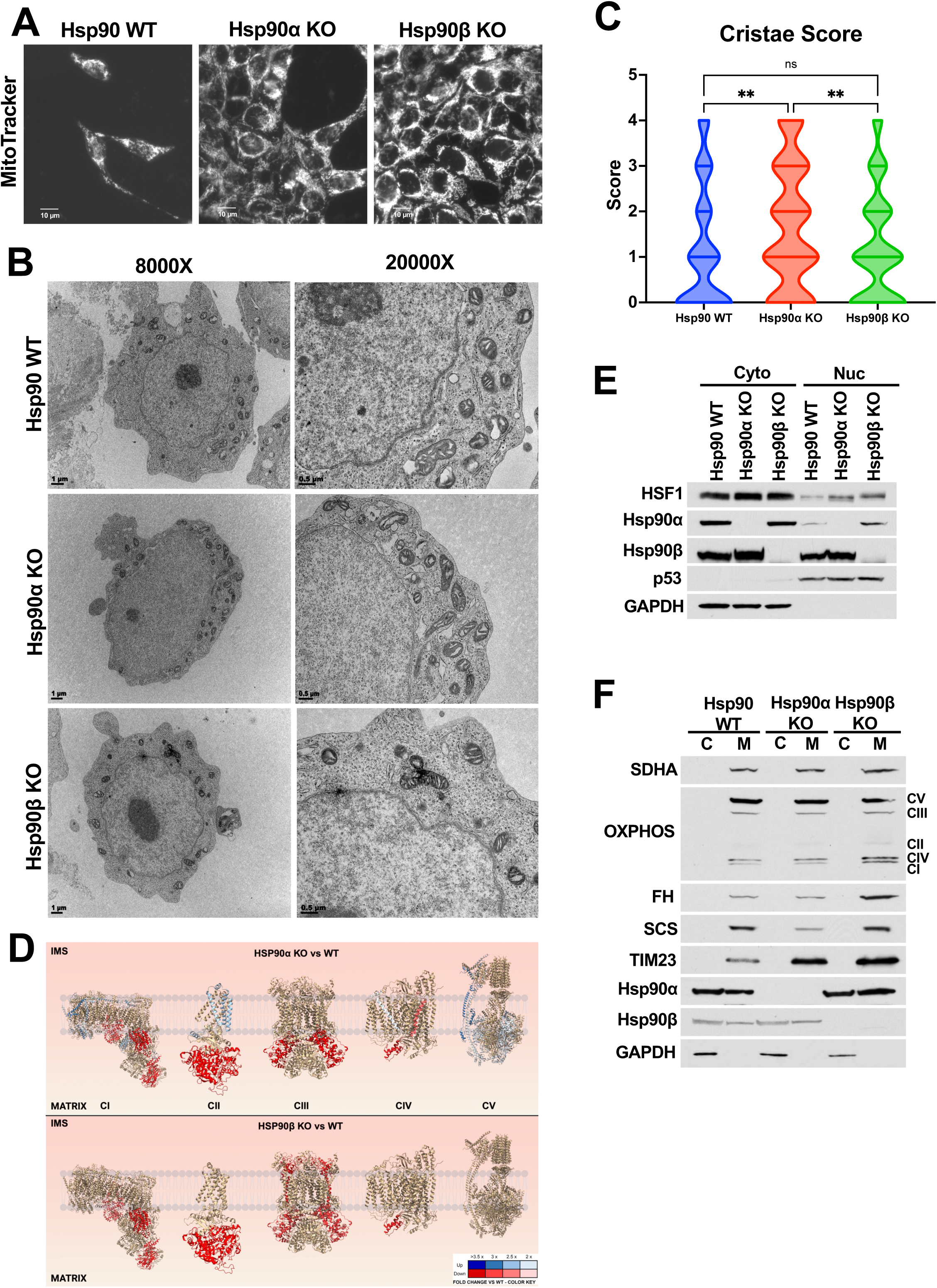
Hsp90s control mitochondrial structure and function. **A)** WT, Hsp90α KO, and Hsp90β KO HEK293 cells were incubated with MitoTracker Green (50 nM, 20 m). Scale bar is 10 µm. **B)** Representative transmission electron microscopy images of WT, Hsp90α KO, and Hsp90β KO HEK293 cells. Scale bar is 1 µm (8000X) or 0.5 µm (20000X). **C)** Mitochondria Cristae Score calculated based on images collected in Figure 1B. **D)** Schematic representation of electron transport chain subunits whose mRNA levels are impacted by loss of Hsp90α or Hsp90β. Data presented as fold change KO/WT. Red subunits indicate decreased mRNA expression, while blue subunits represent increased mRNA expression. PDB IDs: CI: 5xtd; CII: 8gs8; CIII: 5xte; CIV: 5xth, CV: 8h9s. **E)** Cytosolic and nuclear fractions of WT, Hsp90α KO, and Hsp90β KO HEK293 cells were immunoblotted with the indicated antibodies. **F)** Cytosolic and mitochondrial fractions collected from WT, Hsp90α KO, and Hsp90β KO HEK293 cells were immunoblotted with the indicated antibodies.

The electron transport chain (ETC) provides cristae structure^25^. Given the observed variability in cristae organization in the cell models, we first asked whether ETC assembly was impacted following Hsp90 KO. To visualize intact ETC complexes, we performed blue native PAGE. We observed a pronounced reduction in ETC complex formation in Hsp90α KO mitochondria, and a modest reduction in Hsp90β KO mitochondria (**Figure S1E**). We next asked whether the differences we observed in mitochondrial ultrastructure resulted from a failure to transcribe nuclear encoded mitochondrial proteins (**Figure 1D**). Using a respiration-targeted PCR microarray, we found that mRNA levels of many soluble electron transport chain subunits were reduced following loss of Hsp90α or Hsp90β, while many membrane-bound subunits exhibited elevated transcript levels only in Hsp90α KO (**Figure 1D, Table S1**). Based on the inconsistent effects on respiration-associated transcript levels, we tested whether Hsp90 loss impacted transcription factor translocation to the nucleus. We first evaluated heat shock factor-1 (HSF1), as HSF1 activity has been shown to effect transcriptional response downstream of stress caused by loss of Hsp90 function^26,27^, and is implicated in regulating mitochondrial chaperone expression^28^. We observed HSF1 induction in both Hsp90 KO cell lines, but not the control transcription factor p53 (**Figure 1E**) demonstrating that transcription factor activation was not a generalized phenomenon. Therefore, we sought to identify which transcription factors could be responsible for regulating ETC transcription. The ChIP-X Enrichment Analysis Version 3 (ChEA3; https://maayanlab.cloud/chea3/^29^) transcription factor prediction tool identified several transcription factors potentially responsible for regulating the expression of these subunits (**Table S2**). Some of the top hits associated with downregulated mitochondrial protein expression are presumed to be Hsp90 clients^30^, suggesting one potential Hsp90-dependent mechanism for transcriptional tuning of respiration.

Despite these transcriptional changes, we observed only a modest elevation in mitochondrial expression and import of ETC subunits (**Figure 1F, S1F**), and a pronounced induction of the mitochondrial membrane transporter TIM23 (**Figure 1F**), indicative a compensatory proteostatic response. Interestingly, we also observed inconsistent expression of the TCA cycle enzymes fumarase (FH) and succinyl-CoA synthetase (SCS) (**Figure 1F**), hinting that Hsp90 KO cells may experience chronic metabolic dysregulation resulting from loss of Hsp90. Taken together, these data suggest that despite the lack of individual Hsp90 isoforms and the observed perturbations in transcription, the minimal protein chaperoning and trafficking requirements were met.

### Loss of Hsp90α or Hsp90β differentially impacts mitochondrial metabolism

The differences in ETC assembly and TCA cycle enzyme expression prompted us to ask whether these cell lines exhibited similar respiratory output. We first evaluated this by culturing cells in high-glucose medium (4500 mg/L), which reduces dependence on glucose oxidation. In this condition, Hsp90α KO cells exhibited significantly reduced oxygen consumption rate profile (OCR, a surrogate for mitochondrial respiration), while WT and Hsp90β KO cells demonstrated similar basal and maximal OCR (**Figure 2A**). Interestingly, oligomycin treatment reduced WT OCR (but not that of Hsp90β KO) to the levels of Hsp90α KO cells, indicating that this reduced OCR was largely due to compromised ATP-linked respiration (**Figure 2A**). To understand these changes in more detail, we measured the activity of ETC Complex I and II *in vitro*. Interestingly, we observed that WT and Hsp90α KO mitochondria had similar levels of ETC Complex I and II activity, while activity in Hsp90β KO cells was significantly increased (**Figure 2B, C**). We then measured OCR after culturing the cells in low glucose medium (1000 mg/L), which increases dependence on respiration. Compared to WT, Hsp90α KO cells demonstrated reduced respiration, while Hsp90β KO cell OCR was dramatically elevated (**Figure 2D, S2A**). Collectively, we observed that Hsp90β KO cells maintained a relatively elevated respiratory phenotype in both conditions, while WT and Hsp90α KO cell respiration was significantly elevated in the high glucose condition. Importantly, this metabolic phenotype is independent of cell proliferation (**Figure S2B**). These findings indicate significant metabolic flexibility associated with the presence of Hsp90β.

**Figure 2).**
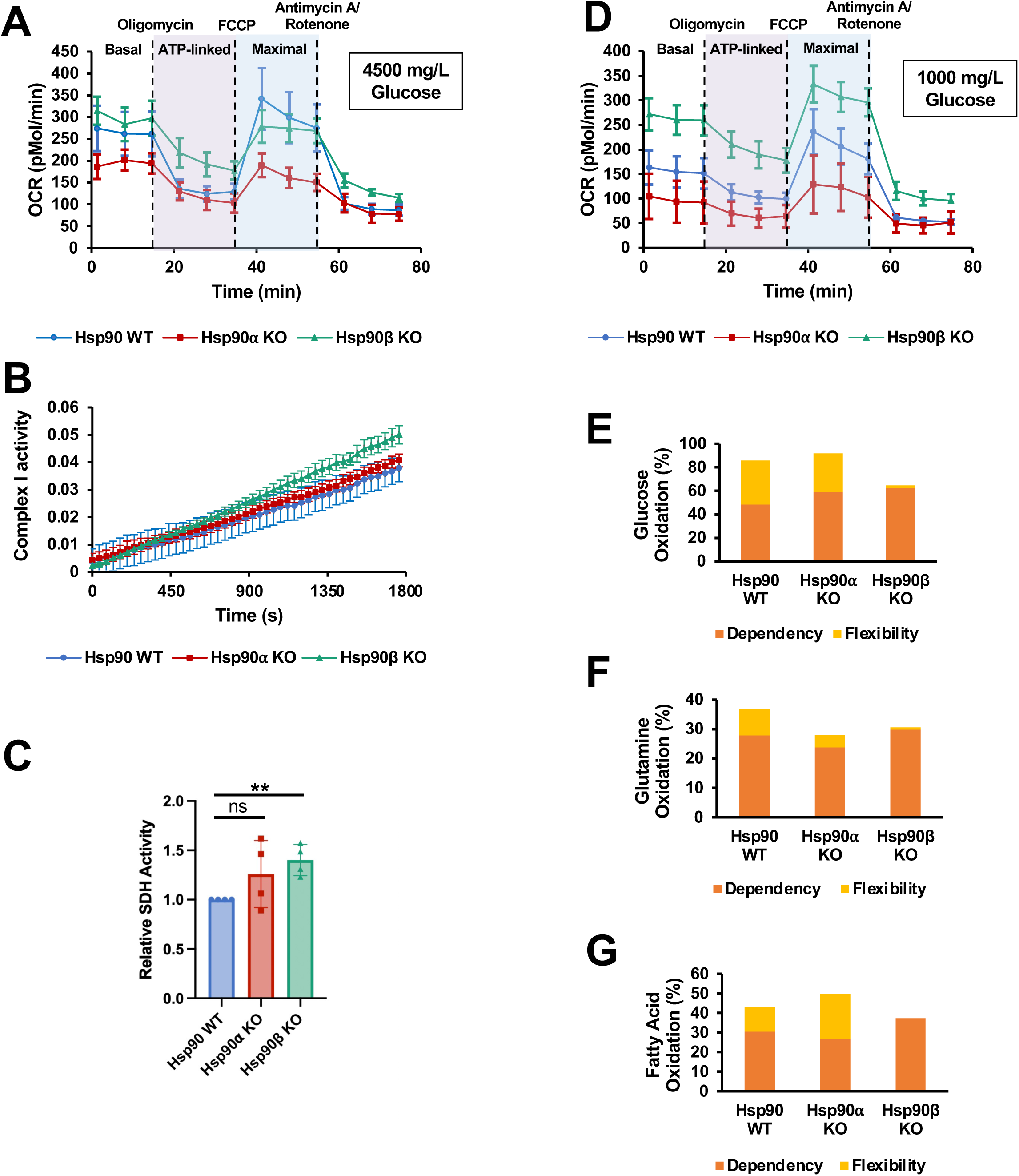
Loss of Hsp90α or Hsp90β differentially impacts mitochondrial metabolism. **A)** Oxygen consumption rate (OCR) based on Mitochondrial Stress Test of WT, Hsp90α KO, and Hsp90β KO HEK293 cells grown in medium containing 4500 mg/L. Error bars represent SD. **B)** Mitochondria isolated from WT, Hsp90α KO, and Hsp90β KO HEK293 cells were assayed for Complex I or **C)** Complex II activity *in vitro*. **D)** Oxygen consumption rate (OCR) measure by Mitochondrial Stress Test of WT, Hsp90α KO, and Hsp90β KO HEK293 cells grown in medium containing 1000 mg/L glucose. Error bars represent SD. **E)** Fuel Flex assay of WT, Hsp90α KO, and Hsp90β KO HEK293 cells grown in medium containing 4500 mg/L glucose. Calculated glucose, **F)** glutamine, **G)** Or fatty acid oxidation dependency or flexibility is shown.

To explore the differences in metabolic flux caused by altered glucose abundance, we performed a Fuel Flex assay, which clearly demonstrated that WT and Hsp90α KO cells exhibited high metabolic flexibility and similar distribution between oxidation of glucose, glutamine, or fatty acid oxidation (**Figure 2E-G, S2C-H**). Hsp90β KO cells demonstrated a marked preference for glucose oxidation with little inherent flexibility (**Figure 2E-G, S2C-H**), suggesting that while the presence of Hsp90α supports respiration, Hsp90β may be regulate metabolic adaptability.

### Hsp90 isoforms regulate diverse aspects of the cellular metabolic network

To begin to understand the mechanism underlying metabolic reprogramming following Hsp90α or Hsp90β loss, we performed targeted metabolomic analysis on whole cell metabolite extracts (**Figure 3A-B, S3A**). Our data showed that the most significantly dysregulated pathways between the WT and Hsp90 KO cells were the pentose phosphate pathway (PPP) and Ala/Asp/Glu metabolism (**Figure 3C, S3B-E**). Hsp90 inhibition has been previously shown to decrease nucleotide biosynthesis in diffuse large B-cell lymphoma cells by reducing the levels of fructose-6-phosphate, ribose-5-phosphate and glyceraldehyde-3-phosphate, among others^31^. Interestingly, constitutive Hsp90 KO did not consistently impact the abundance of these metabolites in our dataset. On the contrary, Hsp90 KO largely corresponded to increased abundance of PPP intermediates (**Figure S3D**). Significantly, most of the identified metabolites relating to Ala/Asp/Glu metabolism were elevated in both Hsp90 KO cell lines, with the exception of pyruvate (**Figure S3E**). We also observed increased levels of several TCA cycle metabolites in Hsp90 KO cells, with Hsp90β KO cells generally exhibiting a greater increase than Hsp90α KO (**Figure S3F**). Importantly, we observed a marked reduction in NAD^+^ levels in both Hsp90 KO cell lines, particularly in Hsp90β KO cells (**Figure S3G**). In agreement, NAD^+^/NADH ratio was significantly decreased in Hsp90 KO cells (**Figure 3D)**. Furthermore, we found minor changes in the levels of NAD^+^-binding proteins in Hsp90 KO cells, and these trends were exacerbated upon testing biotin-NAD^+^ binding capacity (**Figure 3E**). This observation suggests an altered redox environment following loss of Hsp90, potentially contributing to the loss of metabolic flexibility in Hsp90β KO cells.

**Figure 3).**
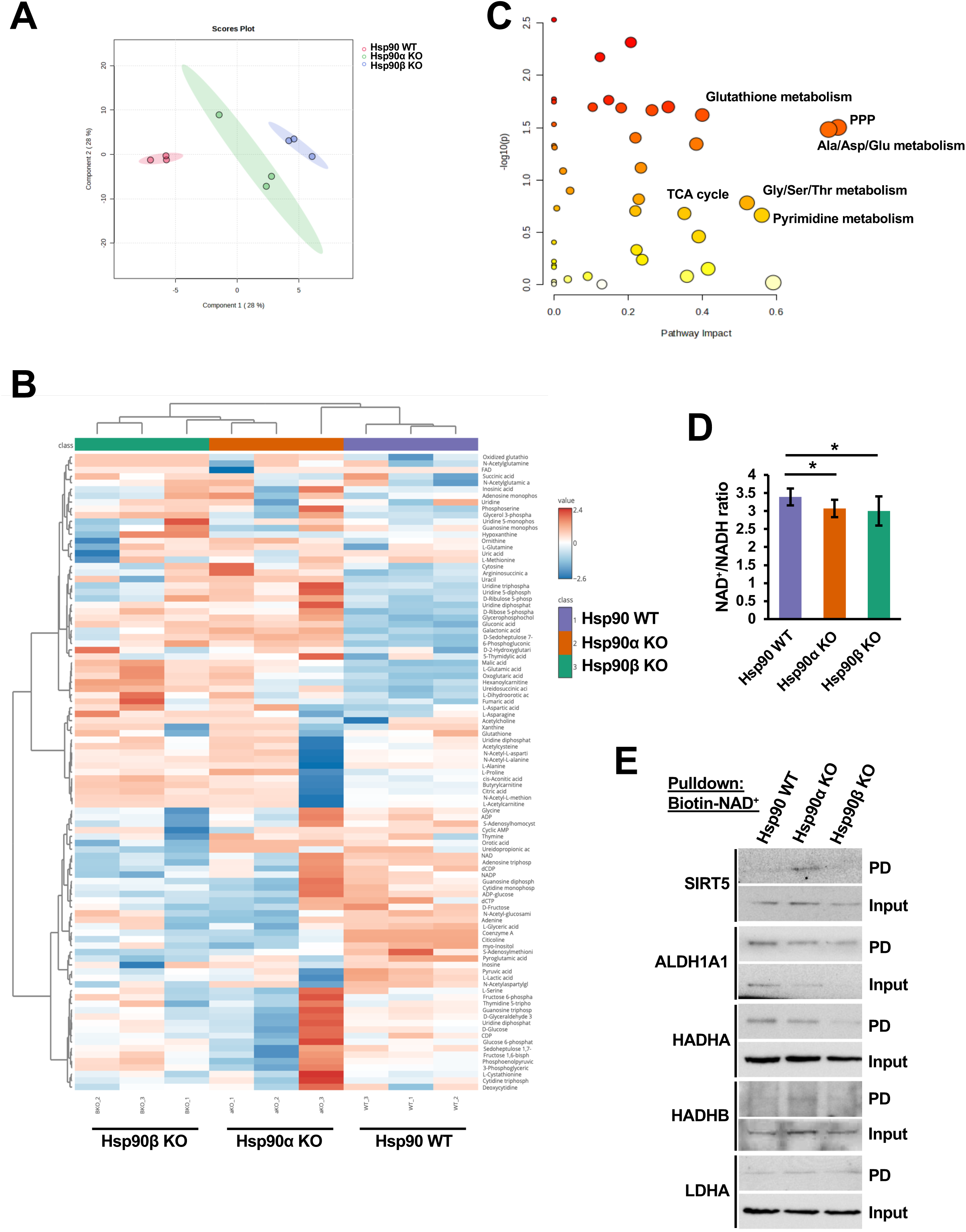
Hsp90 isoforms regulate diverse aspects of the cellular metabolic network. **A)** Principal Component Analysis of WT, Hsp90α KO, and Hsp90β KO HEK293 targeted metabolite composition. **B)** Heat map dendrogram of targeted metabolite abundance across each replicate of WT, Hsp90α KO, and Hsp90β KO HEK293 metabolite extract. **C)** Pathway impact analysis of targeted metabolites isolated from WT, Hsp90α KO, and Hsp90β KO HEK293 cells. **D)** NAD^+^/NADH ratio was calculated based on NAD^+^/NADH-Glo Assay performed in WT, Hsp90α KO, and Hsp90β KO HEK293 cells. **E)** Streptavidin pulldown of biotin-NAD^+^ from WT, Hsp90α KO, and Hsp90β KO HEK293 cell lysate was immunoblotted with the indicated antibodies.

Taken together, these data suggest that the presence of Hsp90α meaningfully promoted glucose oxidation and that individual Hsp90 isoforms control distinct aspects of mitochondrial metabolism.

### The mitochondrial metaboproteome is controlled by Hsp90

To further understand the impact of Hsp90 loss on mitochondrial metabolism, we asked which mitochondrial proteins are clients of mitochondrial Hsp90s. To accomplish this, we Hsp90-binding proteins by immunoprecipitating Hsp90α or Hsp90β from mitochondria of WT HEK293 cells. We observed mitochondrial Hsp90α and Hsp90β interacting with more than 500 intramitochondrial proteins, with some modest preference for individual Hsp90 isoforms (**Figure 4A, Table S3**). Interestingly, Hsp90β alone interacted with several subunits of the ETC complexes, including NDUFA7, NDUFB1, NDUFAB1, NDUFAF5 (CI), SDHD (CII), LYRM7 (CIII), COX18, COX20, MT-CO1 (CIV), and ATP5IF1 (CV), as well as COQ7, involved in ubiquinone biosynthesis (**Figure 4A**). Overall, Hsp90 interactors were significantly enriched for proteins involved in respiration, as well as amino acid and fatty acid metabolism (**Figure 4B, S4A**). Significantly, the majority of ETC subunits were identified in the Hsp90 interactome, including those of the F-type, but not V-type ATPase (**Figure 4C**), as well at least one subunit of each TCA cycle complex (**Figure 4D**).

**Figure 4).**
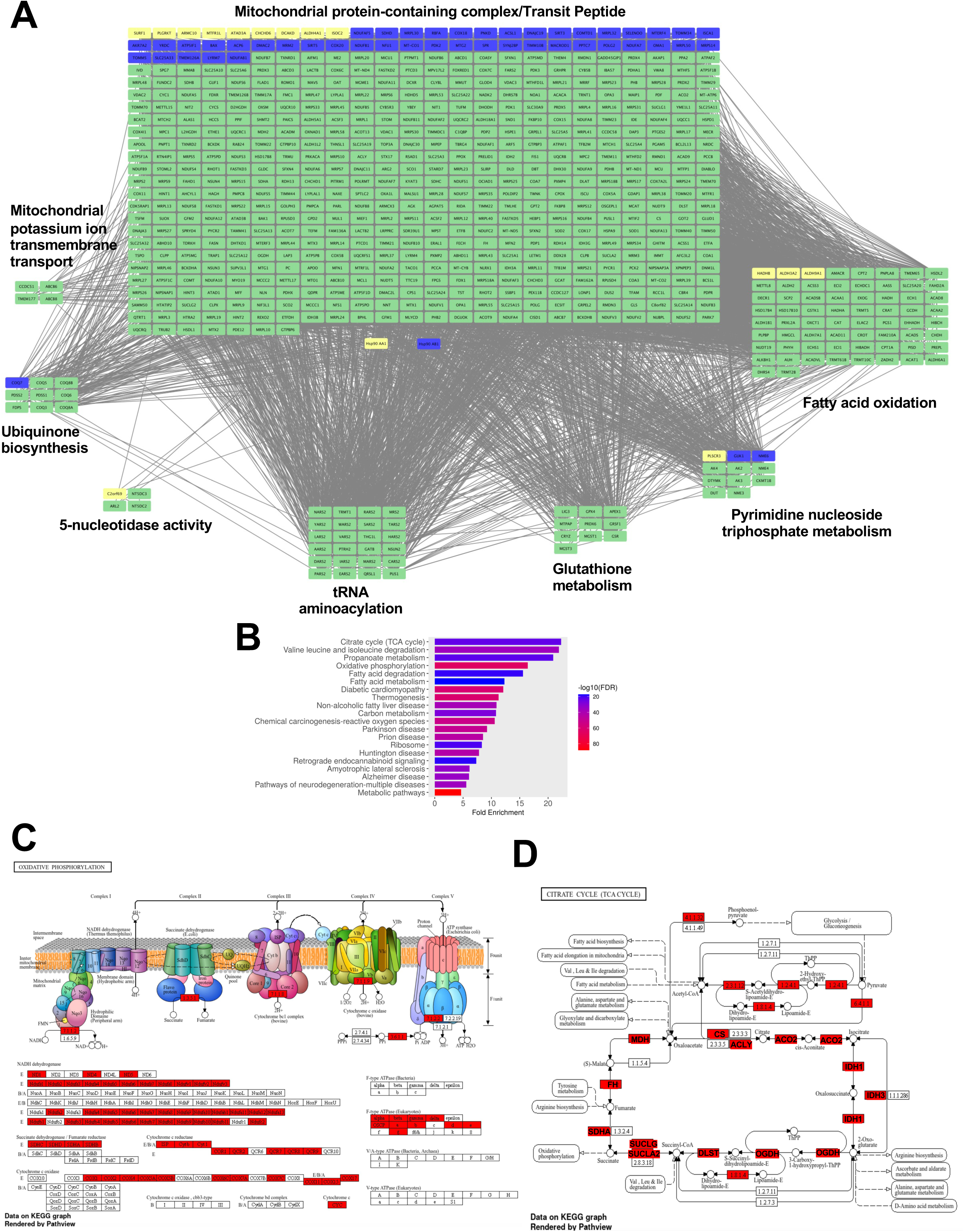
The mitochondrial metaboproteome is controlled by Hsp90. **A)** Protein interaction network of mitochondrial Hsp90α (yellow) and Hsp90β (blue). Proteins found in both interactomes are green. **B)** KEGG enrichment analysis of mitochondrial Hsp90 interacting proteins. **C)** Hsp90 interactors mapped onto a schematic of the ETC or **D)** TCA cycle pathways. Red-labeled proteins were found in the interactome in Figure 1A.

The NAD^+^-dependent deacylases SIRT3/5 were identified as Hsp90β-specific interactors, suggesting a potential mechanism to explain the significantly decreased NAD^+^ levels in Hsp90β KO cells. Similarly, Hsp90α specifically interacted with HADHB and two ALDH family members, integral players in fatty acid oxidation (FAO). In support of this, Hsp90α KO cell OCR responded to the FAO inhibitor etomoxir (ET), in contrast to WT and Hsp90β KO cells, indicative of a potentially important role for Hsp90α in regulating FAO (**Figure S4B**).

### Chaperone compensation supports mitoproteostasis

The mitochondria-dedicated Hsp90 isoform Tumor necrosis factor Receptor-Associated Protein 1 (TRAP1) has an established role in controlling mitochondrial respiration by regulating the activity and assembly of the ETC^17,18^. TRAP1 exhibits distinct structural features from Hsp90α and Hsp90β, suggestive of significant functional divergence, including a significant reduction in the length of the charged N-C linker, and a loss of the Hsp90-MEEVD, a binding motif for interaction with tetratricopeptide repeat-containing proteins^32,33^ (**Figure 5A**). Therefore, we asked whether the three Hsp90 chaperones exhibited unique or overlapping interactors that might explain their respective functions in regulating respiration. To accomplish this, we isolated Hsp90α or Hsp90β from the mitochondria of HEK293 cells and compared their interactors in WT and TRAP1 KO. We initially sought to understand the degree to which TRAP1 loss induced compensatory binding of Hsp90 to mitochondrial proteins. We observed that Hsp90α and Hsp90β both compensate for TRAP1 loss by demonstrating a >2-fold increase in binding to the majority of detected proteins (**Figure S5A-D, Table S4-S7**), with Hsp90α displaying a greater breadth of compensation (78% of interactions increased vs 59%, based on >700 interactions), perhaps due to its stress-responsive nature (**Figure 5B, S5E**). Individually, Hsp90α took on increased interaction with proteins involved in amino acid metabolism and aminoacyl-tRNA biosynthesis (**Figure 5B-C, Table S8**), and acquired *de novo* interactions with proteins regulating nicotinate and nicotinamide metabolism, oxidative phosphorylation, and mitoribosome function (**Figure S5F-G, Table S9**). Compensatory Hsp90β interaction demonstrated a preference for proteins involved in fatty acid biosynthesis and nicotinate and nicotinamide metabolism, while both Hsp90 isoforms demonstrated enrichment for TCA cycle proteins in the absence of TRAP1 (**Figure 5D-E, Table S10**).

**Figure 5).**
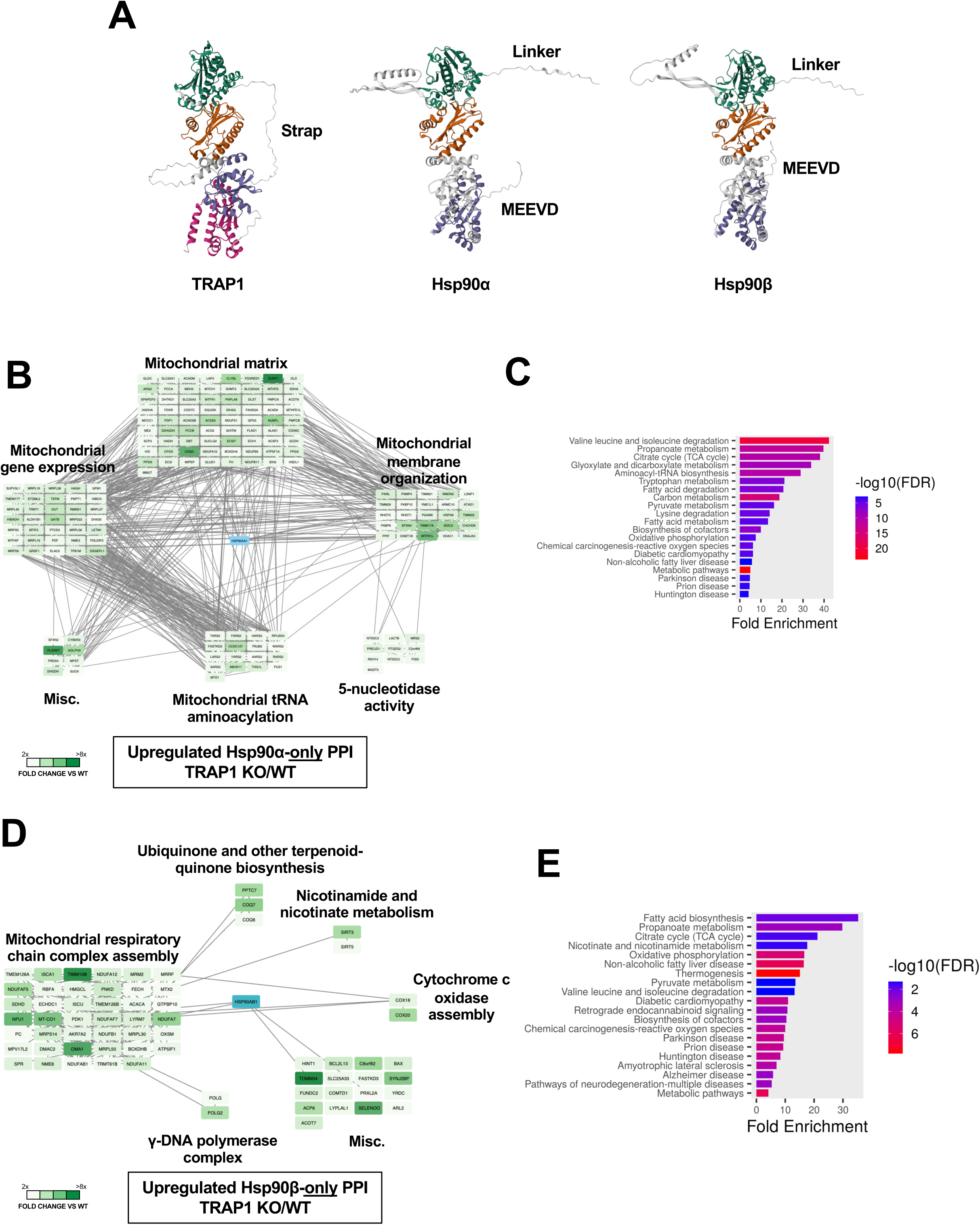
Chaperone compensation supports mitoproteostasis. **A)** Alphafold^55,56^ structure prediction of TRAP1, Hsp90α, and Hsp90β. Colored portions represent predicted discrete structural domains (N-domain: green, Hsp90 large and small middle domains: orange and gray, TRAP1 large and small middle domains: orange and purple, Hsp90 C-domain: purple, TRAP1 C-domain: pink). **B)** Interaction network of proteins whose interaction with Hsp90α only is upregulated in TRAP1 KO HEK293 cells. Increased color intensity corresponds to degree of interaction upregulation. **C)** KEGG enrichment analysis of proteins whose interaction with Hsp90α only is upregulated in TRAP1 KO HEK293 cells. **D)** Interaction network of proteins whose interaction with Hsp90β only is upregulated in TRAP1 KO HEK293 cells. Increased color intensity corresponds to degree of interaction upregulation. **E)** KEGG enrichment analysis of proteins whose interaction with Hsp90β only is upregulated in TRAP1 KO HEK293 cells.

To understand whether Hsp90 chaperone compensation was bidirectional, we next compared TRAP1 interactors in WT, Hsp90α KO, and Hsp90β KO cells. Our data showed that in the absence of Hsp90α or Hsp90β, TRAP1 took on additional interactors previously observed in the Hsp90α or Hsp90β interactomes, suggesting a degree of functional redundancy that was slightly elevated in Hsp90α KO cells (71% of interactions increased vs 60%, based on ∼50 interactors; **Figure S6A**). Interestingly, proteins associated with fatty acid and amino acid metabolism were significantly enriched for TRAP1 protein-protein interactions (PPIs) in Hsp90 KO cells (**Figure S6B-D**), while TCA cycle proteins were significantly underrepresented (**Figure S6E-F, Table S11-17**). We examined a subset of these interactors by immunoblot and observed that differences in PPIs were not due to changes in the abundance or mitochondrial import of chaperone-interacting proteins (**Figure S7A-B**). Overall, we identified an order of magnitude more interactors for Hsp90s compared to TRAP1, resulting in a 14-fold increase in upregulated Hsp90 PPIs in TRAP1 KO compared to TRAP1 PPIs in Hsp90 KO cells (**Figure S7C-D**). Taken together, our data suggests that Hsp90s are significantly more capable of compensating for TRAP1 than the inverse.

### Chaperone compensation supports metabolic flux

We next asked whether Hsp90 isoform compensation manifested in differential regulation of mitochondrial respiration under stress. To test this, we treated WT and TRAP1 KO cells with a sub-lethal dose of the mitochondrial Hsp90 inhibitor Gamitrinib-TPP (G-TPP^34^). Unexpectedly, PCA analysis of the metabolomics data showed that DMSO-treated WT and TRAP1 KO cells clustered more closely to each other than to their isogenic G-TPP-treated counterparts (**Figure 6A-B, S8A**). Furthermore, GTPP-treated WT and TRAP1 KO cells were less similar to each other than their untreated counterparts were, suggesting the presence of a meaningfully interconnected chaperone network disrupted by G-TPP treatment (**Figure 6A, S8A**).

**Figure 6).**
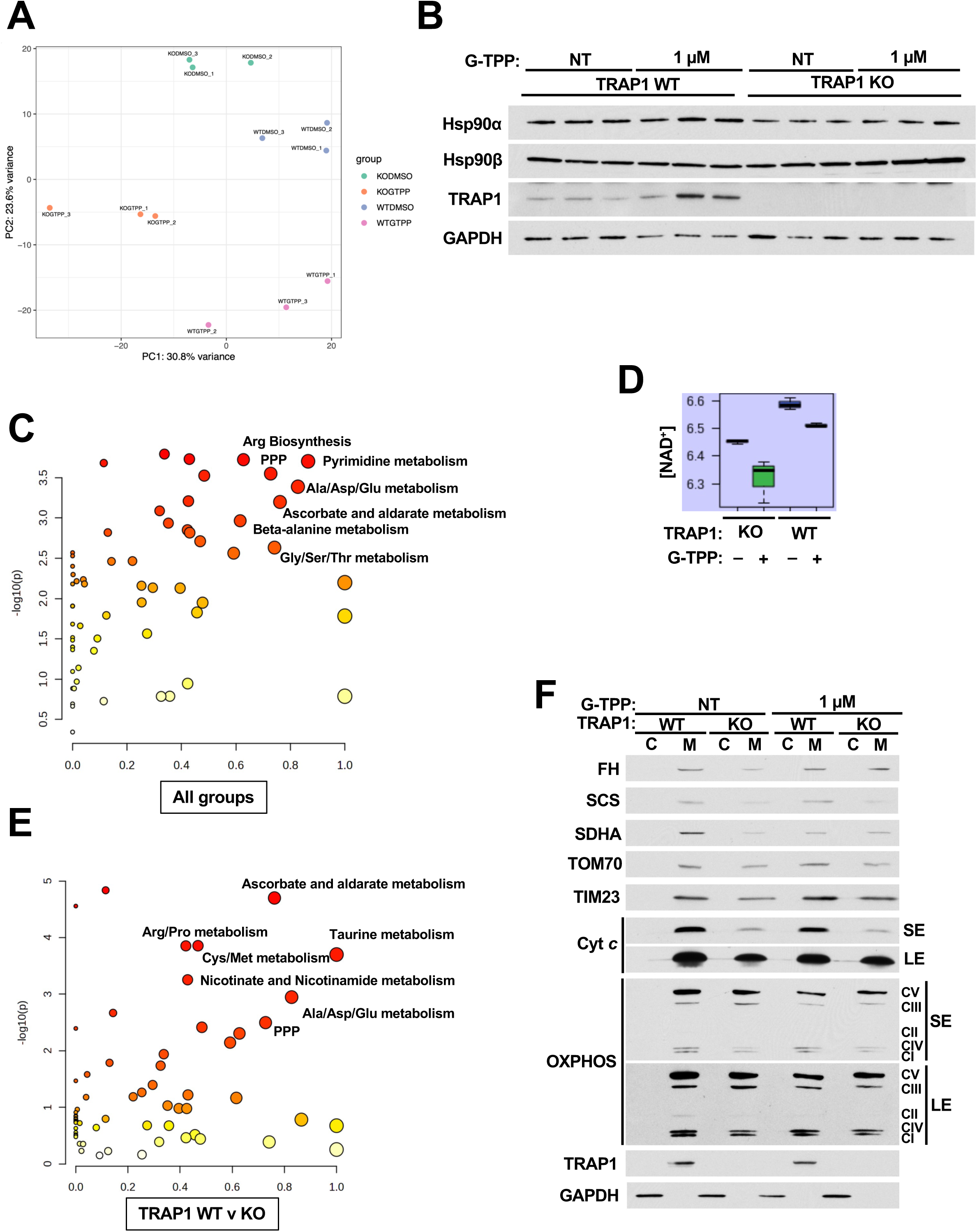
Chaperone compensation supports metabolic flux. **A)** Principal Component Analysis of WT or TRAP1 KO HEK293 cell metabolite composition in the presence or absence of the mitochondrial Hsp90 inhibitor G-TPP **B)** Immunoblot of protein fractions of WT or TRAP1 KO HEK293 cell samples submitted for metabolomics analysis. **C)** Pathway impact analysis of metabolites isolated from WT or TRAP1 KO HEK293 cells in the presence or absence of G-TPP. **D)** NAD^+^ abundance in metabolite extracts from WT or TRAP1 KO HEK293 cells in the presence or absence of G-TPP. **E)** Direct comparison of pathway impact of metabolites isolated from WT or TRAP1 KO HEK293 cells. **F)** Immunoblot of metabolic enzymes from cytosolic or mitochondrial fractions of WT or TRAP1 KO HEK293 cells in the presence or absence of G-TPP. SE - short exposure; LE - long exposure.

We explored differences in chaperone-dependence using Metaboanalyst 6.0. Among all four groups, Ala/Asp/Glu and Gly/Ser/Thr metabolism and PPP function were the primary pathways exhibiting significant alterations (**Figure 6C**). Furthermore, TRAP1 loss reduced cellular NAD^+^ levels, which was exacerbated by mitochondrial Hsp90 inhibitor G-TPP treatment (**Figure 6D**). These observations are consistent with our metabolomic data in Hsp90 KO cells (**Figure 3**), indicating that altered metabolic flux and redox balance is in part dependent on mitochondrial Hsp90 function.

Within the untreated groups, we found that TRAP1 loss was primarily associated with dysregulation of Ala/Asp/Glu, Arg/Pro, and Cys/Met metabolism, as well as ascorbate and taurine metabolism (**Figure 6E**). Interestingly, we observed that TRAP1 loss appreciably decreased UDP-GlcNAc levels (**Figure S8B**), consistent with our prior observations^35^. Pyrimidine metabolism was similarly dysregulated in both cell lines following mitochondrial Hsp90 inhibitor G-TPP treatment (**Figure S8C-D, S9A-F**), indicating that this pathway is Hsp90-dependent, but TRAP1 independent. Interestingly, we found that low dose G-TPP treatment had distinct effects on mitochondrial metabolic protein levels in TRAP1 WT and KO cells (**Figure 6F**), an indication that differential chaperone requirements for these proteins layer significant nuance onto metabolic flux regulation. Taken together, our data strongly support distinct mechanisms by which Hsp90α, Hsp90β and TRAP1 control metabolic flux distribution.

## DISCUSSION

Hsp90 chaperones play a significant role in regulating biological processes through their ability to control protein folding, maturation and activation. To fully understand how Hsp90 chaperone function impacts metabolism, we applied integrated high-resolution proteomic, metabolomic, and transcript-level datasets along with functional assays. We have demonstrated both specific and overlapping roles for three Hsp90 chaperones in controlling metabolite flux and metabolic flexibility. Perhaps most importantly, we have quantitatively identified a set of chaperone-protein interactions and metabolite signatures in response to loss of individual Hsp90 chaperones or inhibition of the mitochondrial Hsp90 chaperone pool.

Cytosolic Hsp90s are responsible for delivering precursor proteins to the translocase of the mitochondrial membrane (TOM) complexes for import^36^. In our individual Hsp90 KO cells, we found inconsistent effects on mitochondrial protein levels, suggesting that a single Hsp90 may be sufficient to support this activity. Our microarray data suggest a second potential compensatory mechanism regulating the transcription of some mitochondrial proteins that may serve to ensure an appropriate amount of these proteins are imported. Furthermore, the impact on mitochondrial structure and electron transport chain complex assembly also indicates a role for Hsp90 in ETC-mediated cristae formation and organization. A similar function has been also shown for TRAP1, however whether these chaperones have mechanistic overlap in this role is unclear. Collectively, our data indicate a multi-faceted role for Hsp90s in regulating all aspects of the mitochondrial protein functional cycle.

Our data clearly demonstrate a role for intra- and extra-mitochondrial Hsp90 function in regulating nucleotide, amino acid, and fatty acid metabolism. In agreement with our work, previous proteomic studies have found Hsp90 interaction with enzymes in glycolysis, the TCA cycle, and one carbon metabolism^37,38^, and a pronounced effect of Hsp90 inhibitors on both glycolysis and respiration in cancer cells^39,40^. However, relative to cytosolic Hsp90 clients, chaperone dependency among mitochondrial proteins is poorly understood. We have described mitochondrial Hsp90 interactors, and categorized their interactions with individual Hsp90 chaperones. Based on our observation of altered chaperone compensation capacity, it is conceivable that Hsp90 and TRAP1 may have different roles with respecting to ‘holding’ or ‘folding’ proteins in the mitochondria, and potentially even function in a sequential manner. Unraveling this interplay will be key to understanding the application and efficacy of isoform-specific Hsp90 inhibitors^41–44^.

Our data implicate Hsp90s as critical regulators of NADH homeostasis through several interrelated mechanisms. Hsp90 or TRAP1 loss disrupts NAD^+^/NADH ratio by reducing NAD^+^ levels and compromises the ability of mitochondrial NAD^+^-binding proteins to interact with NAD^+^. The malate-aspartate shuttle (MAS) is responsible for transporting reducing equivalents in the mitochondria to maintain NADH levels for oxidative phosphorylation^45^. Our data show that GOT2, the mitochondrial isoform of glutamic-oxaloacetic transaminase and linchpin of the MAS, is among Hsp90 and TRAP1 interactors. Furthermore, malate and aspartate levels are upregulated while 2-oxoglutarate levels are decreased, both in Hsp90 and TRAP1 KO cells as well as in response to G-TPP treatment. Interestingly, minimal changes in NADH levels were observed, but NADP^+^ abundance mirrored NAD^+^. These key observations suggest that Hsp90s control MAS function, and potentially also contribute to disrupted redox homeostasis across the cell.

Despite limited organelle-specific resolution of metabolite abundance and Hsp90 function, certain examples remain particularly informative. For example, mitochondrial Hsp90 inhibitor G-TPP treatment significantly reduced orotate formation. The mitochondrial enzyme DHODH catalyzes orotate production and is a significantly increased Hsp90α interactor in TRAP1 KO cells. Notably, this is the only step of pyrimidine biosynthesis that occurs in the mitochondria. This pathway is significantly impacted in both Hsp90 and TRAP1 KO cells, as well as in response to G-TPP treatment, in agreement with previous observation^31^. However, the relative contributions of each chaperone to the folding, maturation, and activation of enzymes in this pathway remains unresolved. Several similarly compelling vignettes can be made for other metabolite-enzyme pairs based on our data, highlighting the potentially significant metabolic dependencies of Hsp90 chaperone activity.

Post-translational modification (PTM) is a major regulatory mechanism for Hsp90 chaperones^46,47^, and can also potentially contribute to localization or retention in specific subcellular compartments. Previous work has shown that nitration of Hsp90-Y33 promotes mitochondrial translocation of Hsp90, with a concomitant negative effect on respiration^20^. Tyrosine nitration is induced by oxidative stress, suggesting a potential feedback mechanism regulating Hsp90 control of oxidative metabolism. A mechanistically similar PTM, nitrosylation, has been observed to regulate TRAP1 activity and stability, with an opposing effect on respiration^48,49^. The ability to simultaneously modulate intramitochondrial Hsp90 isoform function under oxidative conditions may provide cells the ability to fine-tune metabolism under stress. Whether additional PTMs are involved in Hsp90 mitochondrial trafficking or activity is unknown.

Dysregulation of chaperone function has been implicated in various diseases, including cancer^50^, neurodegenerative disorders^51,52^, and cystic fibrosis^53^, where proper chaperone function is critical for preserving protein homeostasis in cells. As such, Hsp90 inhibitors have consistently been observed to have strong anti-proliferative and anti-cancer effects^54^. Our data indicate a substantial impact of Hsp90 loss on nucleotide, amino acid, and fatty acid biosynthesis, suggesting that the decreased proliferation of Hsp90 KO cells we observed results from a lack of essential macromolecules required for cell division. It is therefore conceivable that a critical function of Hsp90 isoforms in cancer is to maintain these anabolic pathways. Understanding the individual roles of these chaperones in the regulation of disease-associated pathways will help to refine therapies based on targeting specific chaperone isoforms.

## LIMITATIONS

The current study has three noteworthy limitations that are essential to acknowledge when interpreting the data presented here. First, the use of constitutive Hsp90 isoform knockout cells to carry out the experiments, allows for the possibility of cellular adaptation that may not faithfully phenocopy the effects of acute Hsp90 inhibition. Secondly, proteomics data presented in this study have been performed in the absence of protease treatment, which would remove proteins associated with the outer mitochondrial membrane. Therefore, some observed Hsp90 interactions are likely to be the product of Hsp90 associated with mitochondrial protein import complexes. Finally, isoform-specific small molecule Hsp90 inhibitors with mitochondrial tropism are not currently available, preventing direct evaluation of cells containing a single functional Hsp90 isoform within the mitochondria.

**Figure S1) Related to Figure 1**.

A) WT, Hsp90α KO, and Hsp90β KO HEK293 cells were incubated with TMRM (50 nM, 20 m). Scale bar is 25 µm.

**B)** Mitochondria/cell in WT, Hsp90α KO, and Hsp90β KO HEK293 cells based on transmission electron microscopy images (**Figure 1B**).

**C)** Average cristae score per cell based on **Figure 1C**.

**D)** Enlarged images of representative mitochondria from WT, Hsp90α KO, and Hsp90β KO HEK293 cells.

**E)** Blue native PAGE (BN-PAGE) or mitochondria isolated from WT, Hsp90α KO, and Hsp90β KO HEK293 cells. Notable ETC complexes have been indicated by labels.

**F)** Whole cell lysates from WT, Hsp90α KO, and Hsp90β KO HEK293 cells immunoblotted with the indicated antibodies.

**Figure S2) Related to Figure 2**.

**A)** Fold change OCR 4500/1000 mg/L glucose. Related to **Figure 2A-B**.

**B)** Proliferation rate of WT, Hsp90α KO, and Hsp90β KO HEK293 cells grown in medium containing either 4500 or 1000 mg/L glucose measured by WST Proliferation Assay.

**C-H)** Fuel Flex assay OCR measurements over time. Used to calculate **Figure 2E-G**.

**Figure S3) Related to Figure 3**.

**A)** Immunoblot of protein fractions of WT, Hsp90α KO, and Hsp90β KO HEK293 cell samples submitted for targeted metabolomic analysis.

**B)** Schematic of interplay between metabolic pathways. Created with Biorender.com

**C)** Color code of metabolite abundance graphs in **Figure S3C-F**.

**D)** Impact of Hsp90α or Hsp90β KO on pentose phosphate pathway (PPP),

**E)** Ala/Asp/Glu metabolism, or

**F)** TCA cycle flux in HEK293 cells. Color intensity gradient (yellow: low - red: high) corresponds to weighted impact of metabolite abundance on pathway significance.

**G)** NAD^+^ abundance in metabolite extracts from WT, Hsp90α KO, and Hsp90β KO HEK293 cells.

**Figure S4) Related to Figure 4**.

**A)** KEGG term enrichment analysis of “mitochondrial protein-containing complex” interaction cluster from **Figure 4A**.

**B)** Fatty acid oxidation-dependent OCR in WT, Hsp90α KO, and Hsp90β KO HEK293 cells grown in medium containing 4500 mg/L glucose.

**Figure S5) Related to Figure 5**.

**A)** Interaction network of proteins whose interaction with Hsp90α is upregulated in TRAP1 KO HEK293 cells. Increased color intensity corresponds to degree of interaction upregulation.

**B)** KEGG enrichment analysis of proteins whose interaction with Hsp90α is upregulated in TRAP1 KO HEK293 cells.

**C)** Interaction network of proteins whose interaction with Hsp90β is upregulated in TRAP1 KO HEK293 cells. Increased color intensity corresponds to degree of interaction upregulation.

**D)** KEGG enrichment analysis of proteins whose interaction with Hsp90β is upregulated in TRAP1 KO HEK293 cells.

**E)** Percent compensation for Hsp90s in TRAP1 KO cells.

**F)** Interaction network of proteins exhibiting *de novo* Hsp90α interaction in TRAP1 KO cells.

**G)** KEGG enrichment analysis of proteins exhibiting *de novo* Hsp90α interaction in TRAP1 KO cells.

**Figure S6) Related to Figure 5**.

**A)** Percent compensation by TRAP1 in Hsp90 KO cells.

**B)** KEGG enrichment analysis of proteins whose interaction with TRAP1 is upregulated in Hsp90α KO HEK293 cells.

**C)** KEGG enrichment analysis of proteins whose interaction with TRAP1 is upregulated only in Hsp90α KO HEK293 cells.

**D)** KEGG enrichment analysis of proteins whose interaction with TRAP1 is upregulated in Hsp90β KO HEK293 cells.

**E)** KEGG enrichment analysis of proteins whose interaction with TRAP1 is downregulated only in Hsp90α KO HEK293 cells.

**F)** KEGG enrichment analysis of proteins whose interaction with TRAP1 is downregulated only in Hsp90β KO HEK293 cells.

**Figure S7) Related to Figure 5**.

**A)** Levels of a subset of Hsp90- or TRAP1-interacting proteins immunoblotted in whole cell lysates from individual Hsp90 isoform KO cell lines.

**B)** Levels of a subset of Hsp90- or TRAP1-interacting proteins by immunoblotted in cytosolic or mitochondrial fractions of individual Hsp90 isoform KO cell lines.

**C)** Relative number of upregulated chaperone interactors in chaperone KO cell lines.

**D)** Chaperone compensation score of Hsp90 isoforms.

**Figure S8) Related to Figure 6**.

**A)** Heat map of individual metabolite abundance across each replicate of metabolite extracts from WT or TRAP1 KO HEK293 cells in the presence or absence of G-TPP.

**B)** UDP-GlcNAc abundance in metabolite extracts from WT or TRAP1 KO HEK293 cells.

**C)** Direct comparison of pathway impact of metabolites isolated from WT HEK293 cells in the presence or absence of G-TPP.

**D)** Direct comparison of pathway impact of metabolites isolated from TRAP1 KO HEK293 cells in the presence or absence of G-TPP.

**Figure S9) Related to Figure 6**.

**A)** Color code of metabolite abundance graphs in **Figure 6C**.

**B)** Impact of TRAP1 and G-TPP treatment on pentose phosphate pathway (PPP),

**C)** TCA cycle,

**D)** Pyrimidine metabolism,

**E)** Arg biosynthesis,

**F)** Or Ala/Asp/Glu metabolism TCA cycle flux in HEK293 cells. Color intensity gradient (yellow: low - red: high) corresponds to weighted impact of metabolite abundance on pathway significance.

## Materials and Methods

### Mammalian Cell Culture

HEK293 Hsp90 KO cells were a generous gift of Didier Picard (University of Lausanne) and were reported in^57^. HEK293 TRAP1 KO were a graciously provided by Len Neckers (National Cancer Institute) and were reported in^58^. Cells were cultured in Dulbecco’s Modified Eagle Medium (DMEM; Sigma-Aldrich) supplemented with 10% fetal bovine serum (FBS; Gibco) at 37 °C in a CellQ incubator at 5% CO2.

### Subcellular fractionation

Cells were placed on ice and washed once with ice cold PBS (Millipore-Sigma). Following thorough aspiration to remove PBS, 100 μl of fresh Isotonic Buffer (250 mM Sucrose, 10 mM Tris– HCl pH 7.4, cOmplete protease inhibitor cocktail (Millipore-Sigma), and PhosSTOP (Millipore-Sigma)) was added and cells were scraped and collected in a 1.7 ml Eppendorf tube. The cell suspension was then transferred to a cold dounce homogenizer and disrupted with 10 strokes of a tight-fitting pestle (Ace Glass). Homogenate was transferred back to a 1.7 ml Eppendorf tube and centrifuged at 600 × g for 10 min to pellet cell debris. Supernatant was transferred to a fresh 1.7 ml Eppendorf tube and centrifuged at 10,000 × g for 10 min to pellet mitochondria. Mitochondrial pellet was gently re-suspended in 50 μl fresh Isotonic Buffer for downstream applications.

### Protein extraction and immunoblotting

Cultured cells were placed on ice and washed once with ice cold PBS (Millipore-Sigma). Following thorough aspiration to remove PBS, 200 μl of fresh lysis buffer (20 mM Tris–HCl (pH 7.4), 100 mM NaCl, 1 mM MgCl2, 0.1% NP40, cOmplete protease inhibitor cocktail (Millipore-Sigma), and PhosSTOP (Millipore-Sigma)) was added to the plate and cells were scraped from the plate and collected in a 1.7 ml Eppendorf tube. The cell suspension was sonicated on ice for 2 s using a probe sonicator (QSonica) and centrifuged for 7 min at 15,000 × g to pellet cell debris. The supernatant containing the soluble protein was transferred to a clean 1.7 ml Eppendorf tube and quantified using a Bradford assay (Bio-Rad) according to the manufacturer’s protocol. Nitrocellulose membranes were immunoblotted with primary antibodies (**Table S18**) followed by the indicated HRP-conjugated secondary antibodies (Cell Signaling Technology).

### Immunoprecipitations and Pulldowns

For endogenous immunoprecipitation, protein lysates or isolated mitochondria were incubated with the indicated antibodies overnight at 4 °C with rotation followed by Protein G agarose (ThermoFisher Scientific) for an additional 1 h at 4 °C with rotation. For biotinylated pulldowns, protein lysates were incubated with indicated concentrations of biotinylated-NAD^+^ (R&D Systems) for 1 h at 4 °C with rotation followed by streptavidin-agarose beads (ThermoFisher Scientific) for 1 additional h at 4 °C with rotation. Agarose pellets were washed 4 times with fresh lysis buffer with vortexing and eluted at 95 °C in 5x Laemmli buffer. Precipitated proteins were separated by SDS-PAGE and transferred to nitrocellulose membranes for immunoblotting or submitted for mass spectrometry analysis.

### Drug Treatment

WT and TRAP1 KO HEK293 cells were treated with 1 µM Gamitrinib-TPP (G-TPP; MedchemExpress) for 18 h prior to subcellular fractionation or metabolite extraction.

### Fluorescence Microscopy

WT, Hsp90α KO, and Hsp90β KO HEK293 were incubated with 50 nM of MitoTracker (Invitrogen) or tetramethylrhodamine, methyl ester (TMRM; Invitrogen) for 20 m at 37 °C. Following incubation, medium was removed and replaced with pre-warmed DMEM + 10% FBS and cells were imaged immediately on an ECHO Revolve R4 Microscopy System (Discover Echo Inc.).

### Cristae Scoring

Mitochondria cristae analysis was conducted using ImageJ as previously defined in Lam *et al.* ^24^. Mitochondria were quantified and cristae scores were assigned based on cristae abundance and shape. Scores were determined in accordance with published criteria: 0 for no distinct crista, 1 for >50% of area without cristae, 2 for >25% of area without cristae, 3 for >75% of area with cristae, but irregularly shaped, 4 for many regular cristae in mitochondria area.

### Transmission Electron Microscopy

Cells were trypsinized, resuspended in PBS, collected in tubes and centrifuged at 300 x g for 10 m to pellet. Cell pellets were washed 2 additional times with PBS for 5 m each with centrifugation (300 x g/10 m). Supernatant was removed and replaced with fixative (2.5% Glutaraldehyde + 2.0% Paraformaldehyde in PBS – Electron Microscopy Sciences (EMS), Hatfield PA), and then the cell pellet was resuspended and tubes placed on ice and allowed to fix for 2 h. Cells were then washed 3x with PBS for 5 m each with centrifugation (300 x g/10 m), enrobed in 4% agarose, and post-fixed for 1.5 h with 1% Osmium Tetroxide in PBS on ice (EMS, Hatfield PA). Cells were then washed 4 times with PBS for 5 m each, placed in a fresh vial containing distilled water, and dehydrated through a graded ethanol series to 100%, after which propylene oxide was exchanged 2 times for 15 m each. Following manufacturers standard protocol, cells were infiltrated with and embedded in Embed-812 epoxy resin (EMS, Hatfield PA). Post polymerization, sample blocks were trimmed to expose the cell pellet, then sectioned at 70 nm with a UC7 ultramicrotome (Leica Microsystems, Wetzler Germany) equipped with a diamond knife (Diatome, Quakertown PA). Sections were collected onto 200 mesh grids, allowed to dry, then counter stained with UranyLess for 4 m and lead citrate for 3 m (EMS, Hatfield PA). Cells were then observed with a JEM-1400 Transmission Electron Microscope (TEM) operated at 80kV (JEOL USA Inc., Peabody MA) and digital images acquired with an Orius SC-1000 CCD camera (Gatan Inc., Pleasanton CA) in the Upstate TEM Core.

### Blue Native PAGE

Mitochondria were isolated as described above. 25 µg of mitochondria were loaded onto the gel, then run and stained using the Invitrogen BN-PAGE system, as previously reported^59^.

### PCR Microarray

RNA was isolated from cells using the RNA Isolation kit (Ambion) according to the manufacturer’s protocol. iScript cDNA Synthesis kit was used to generate cDNA according to manufacturer’s protocol. RT^2^ Profiler PCR Array Human Mitochondrial Energy Metabolism Pathway Plus (PAHS-008Y; Qiagen) was performed according to manufacturer’s protocol using 20 ng RNA-equivalent of cDNA. Significant alterations were color-coded according to the following: Positive fold change is colored blue, and downregulation is colored red. Dark blue signifies a fold change exceeding 3.5-fold, while light blue represents a fold change between 2.0 and 2.49-fold. Conversely, gene downregulation is illustrated by the red-colored chains. Similar to upregulation, the shade of red signifies the fold change. Dark red denotes a fold change greater than 3.5, while light red indicates a fold change between 2.0 and 2.49.

### WST Proliferation Assay

Cells were plated at 10^5^ cells/well. After 72 h of culture, MTT assay was performed as described in the Quick Cell Proliferation Colorimetric Assay Kit Plus (BioVision).

### Complex II/Succinate dehydrogenase Activity Assay

SDH activity was measured in 25 µg of mitochondrial lysate using the Colorimetric Succinate Dehydrogenase *in vitro* activity assay (abcam), according to the manufacturer’s protocol.

### Complex I Activity Assay

Complex I activity was measured in 25 µg of mitochondrial lysate using the Colorimetric Complex I *in vitro* activity assay (abcam), according to the manufacturer’s protocol.

### Metabolic flux analysis

Cells were seeded at 5 x 10^5^/well in a 96-well assay plate. Following overnight incubation, cells were treated or subject to media changes as indicated for 16 h prior to assay initiation. The indicated assay was then carried out on an Agilent Seahorse XFe96 instrument according to the manufacturer’s protocol.

### Metabolomic Analysis

For untargeted metabolomics, cells were cultured in indicated medium for 24 h prior to metabolite extraction. For isotope tracing targeted metabolomics, cells were cultured in low-glucose DMEM supplemented with 3.5 g/L ^13^C6 glucose for 30 m prior to metabolite collection. Metabolites were methanol-extracted from cells at 80% confluency. Briefly, cells were washed one time with PBS on wet ice. After thorough aspiration, cells were moved to dry ice and scraped in 700 µl ice cold 80% methanol. Extract was centrifuged at 14000 x g for 20 m. Supernatant was transferred to a new tube and samples were completely dried using a vacufuge. Mass spectrometric analyses were performed at the Weill Cornell Proteomics and Metabolomics Core Facility. Metabolomic data was log transformed and normalized by Pareto scaling followed by visualization using MetaboAnalyst 6.0 web-based tools.

### Proteomic Analysis

The indicated proteins were immunoprecipitated from Hsp90 WT, Hsp90α KO, Hsp90β KO, TRAP1 WT and TRAP1 KO HEK293 cells in duplicate. Mass spectrometric analyses were performed at the Weill Cornell Proteomics and Metabolomics Core Facility. The identified proteins were cleaned using the following criteria: Greater than one unique peptide and present in a minimum of two replicates. The proteins were normalized to the intensities of the precipitated proteins (TRAP1, Hsp90α or Hsp90β) and averaged across replicates. Mitochondrial protein interactors were identified by cross-referencing to the MitoCarta 3.0 database (Broad Institute). Significant fold changes are colored green (greater than 1.9) or red (less than 0.5). Cytoscape v3.10.3 was used to create interactomes based on STRING PPI networks (STRING, version 12.0). ShinyGO v0.80 was used to perform Go/KEGG analyses.

### Chaperone Compensation Calculation

Percent chaperone compensation was calculated using the following formula: ((Upregulated PPIs-downregulated PPIs)/Total PPIs)*100. Chaperone compensation score was calculated using the following formula: Upregulated Hsp90 PPI in TRAP1 KO/Upregulated TRAP1 PPI in Hsp90 KO.

### NAD/NADH-Glo Assay

Cells were seeded at 10^6^/well in a clear tissue-culture treated 96-well plate. After 24 h, the assay was performed in a white 96-well plate according the manufacturer’s protocol (Promega) and read on a Tecan Spark plate reader.

### Quantification and Statistical Analysis

The data presented are the representative of three biological replicates, unless otherwise specified. All statistics were performed using GraphPad Prism version 7.00 for Windows (GraphPad Software, La Jolla, California, USA, https://www.graphpad.com). Statistical significance was ascertained between individual samples using a parametric unpaired t-test. Significance was denoted as asterisks in each figure: *P < 0.05; **P < 0.01; ***P < 0.001; ****P < 0.0001. Error bars represent the standard deviation (SD) for three independent experiments, unless otherwise indicated.

### Data and Software Availability

The mass spectrometry proteomics data have been deposited to the ProteomeXchange Consortium via the PRIDE^60^ partner repository with the dataset identifier PXD069999. Metabolomics datasets have been uploaded to Metabolomics Workbench.

## Acknowledgements

The authors thank Mehdi Mollapour, Dimitra Bourboulia, and Gennady Bratslavsky for support, mentorship, and stimulating scientific discussions. The authors thank Benjamin Zink, manager of the Upstate Medical University Transmission Electron Microscopy Core for TEM imaging services, and Xiaowen Wang for technical assistance. The authors thank Guoan Zhang, director of the Weill Cornell Medicine Proteomics and Metabolomics Core for mass spectrometry analysis and invaluable input.

## Author Contributions

Conceptualization, M.R.W.; Methodology, R.F.P., G.L.M., V.Z.R., L.D.A., S.J.B., M.R.W.; Investigation, R.F.P., G.L.M., J.K.B., V.Z.R., L.D.A., S.J.B., M.R.W.; Writing – Original Draft, M.R.W.; Writing – Reviewing and Editing R.F.P., G.L.M., S.J.B., J.R.W., M.R.W.; Funding Acquisition, M.R.W.; Resources, M.R.W.; Supervision, J.R.W., M.R.W.

