## Supplementary figures and images for "Coordinated regulation of the metaboproteome by Hsp90 chaperones controls metabolic plasticity"

### Supplemental Figures

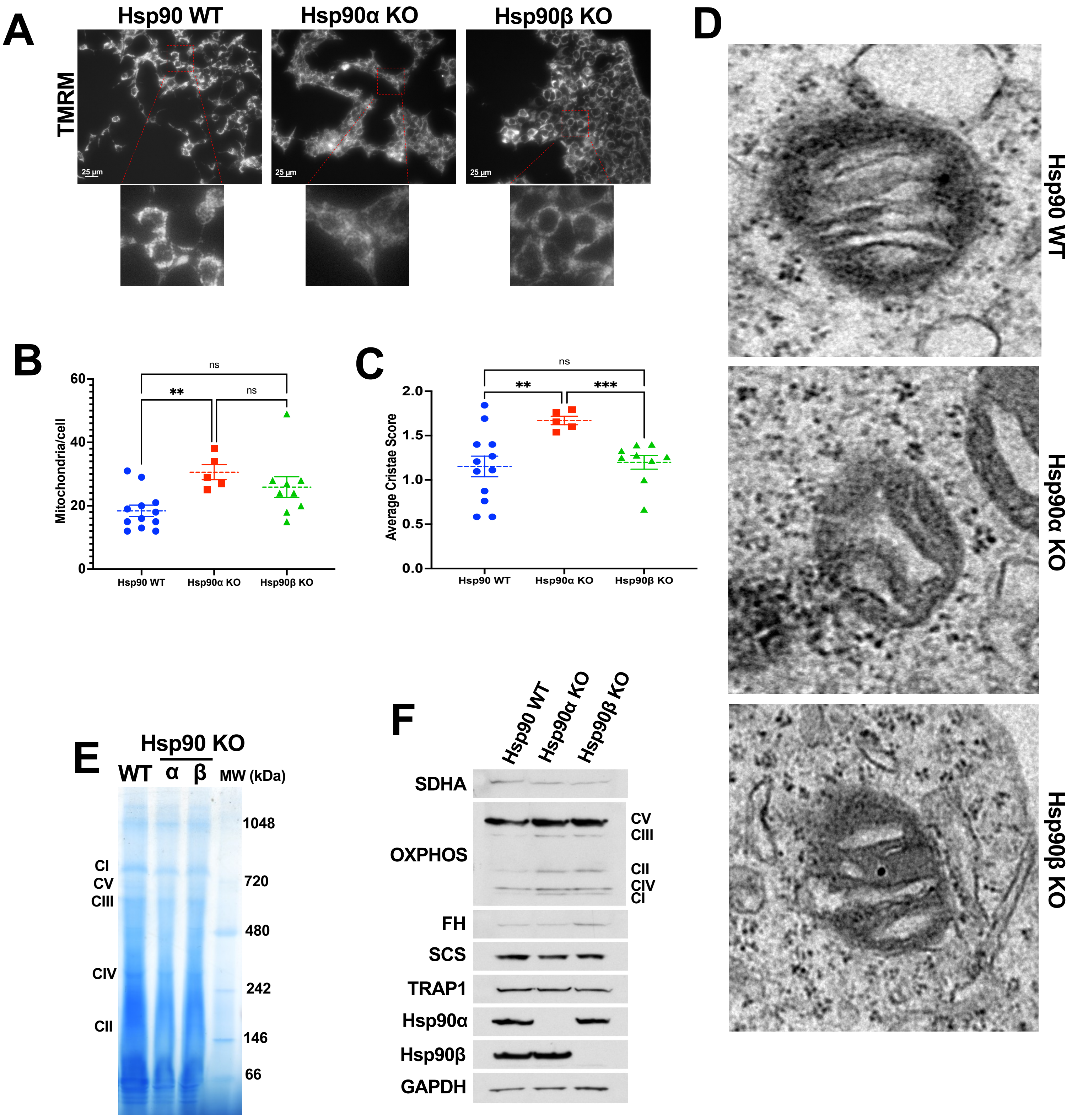

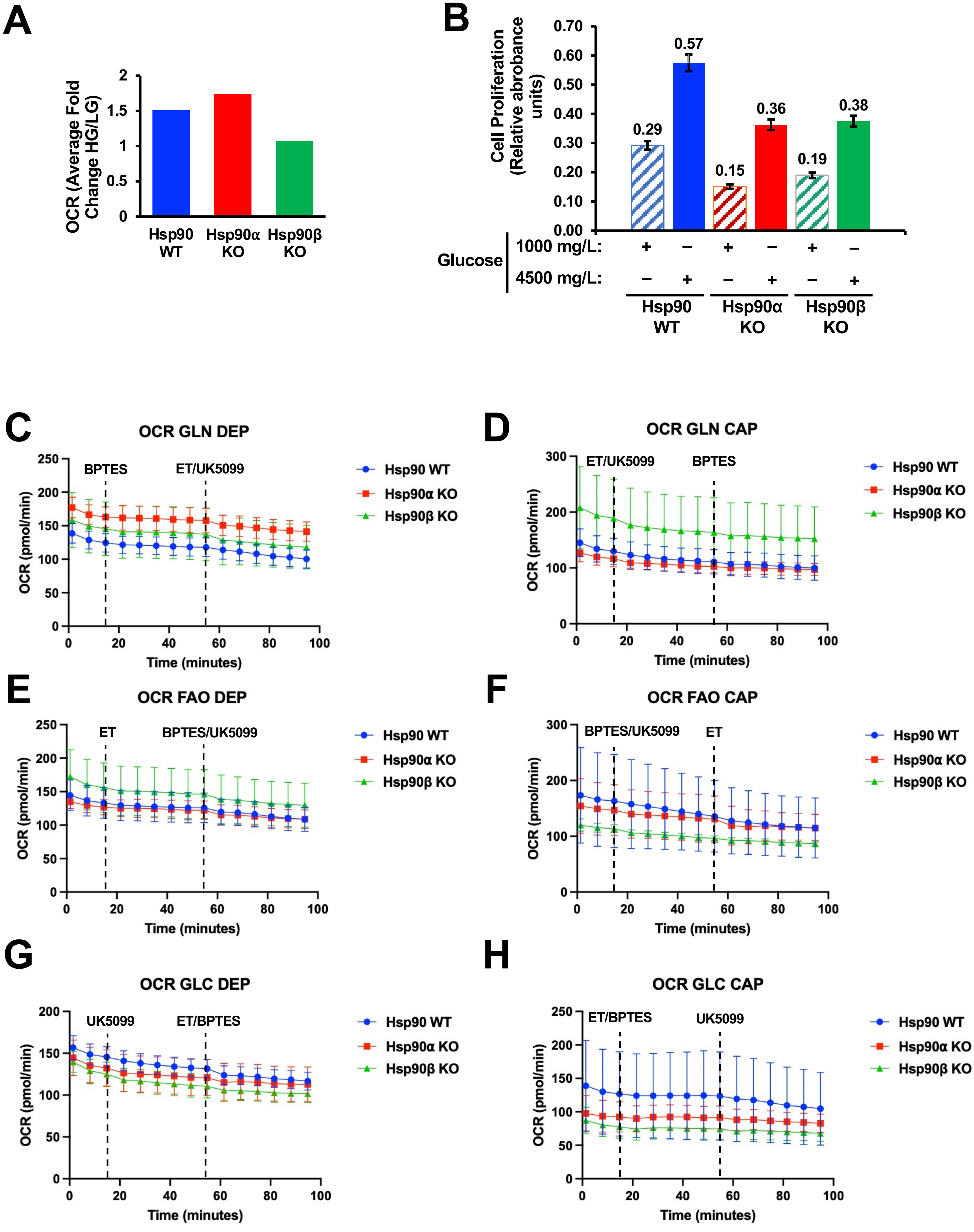

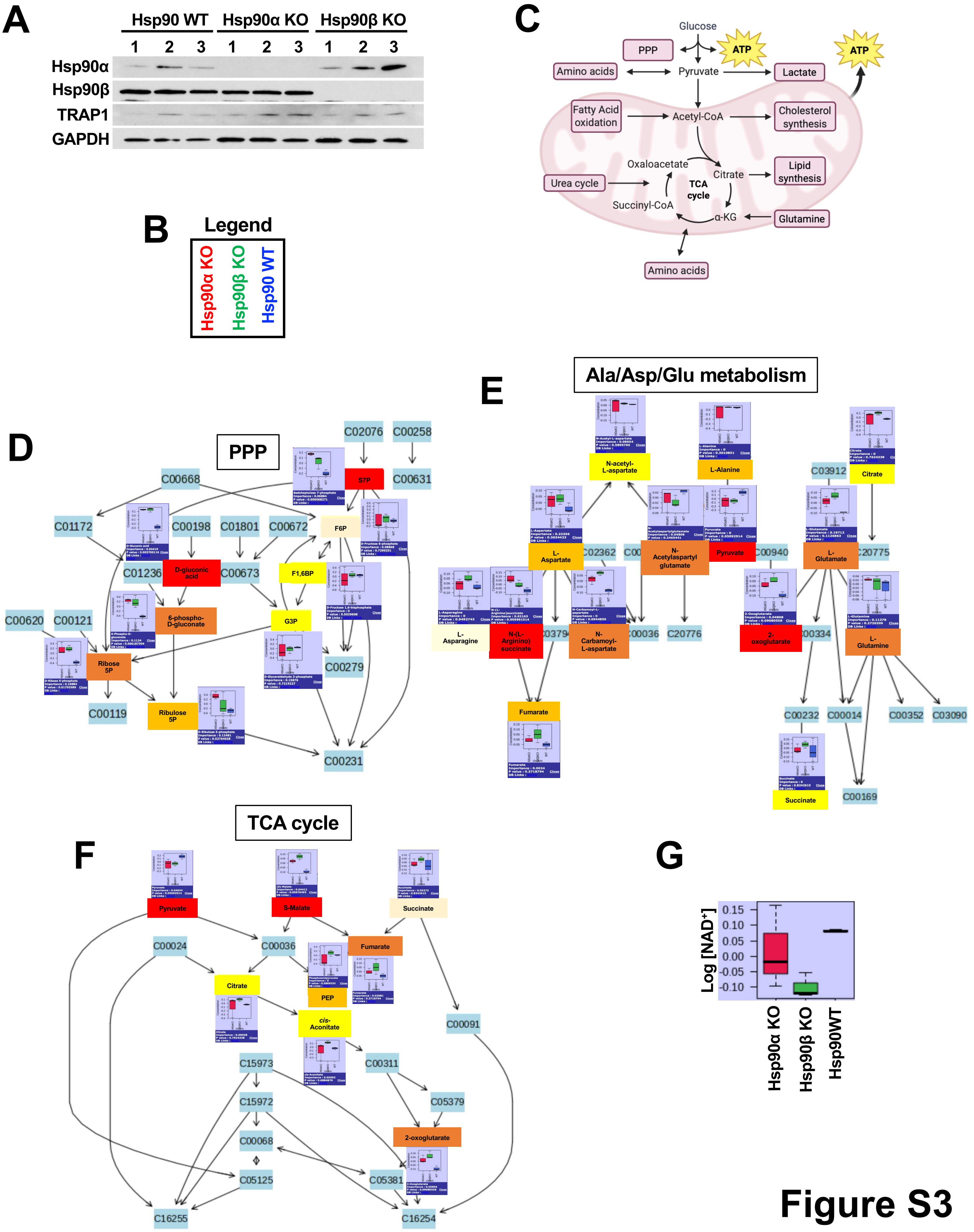

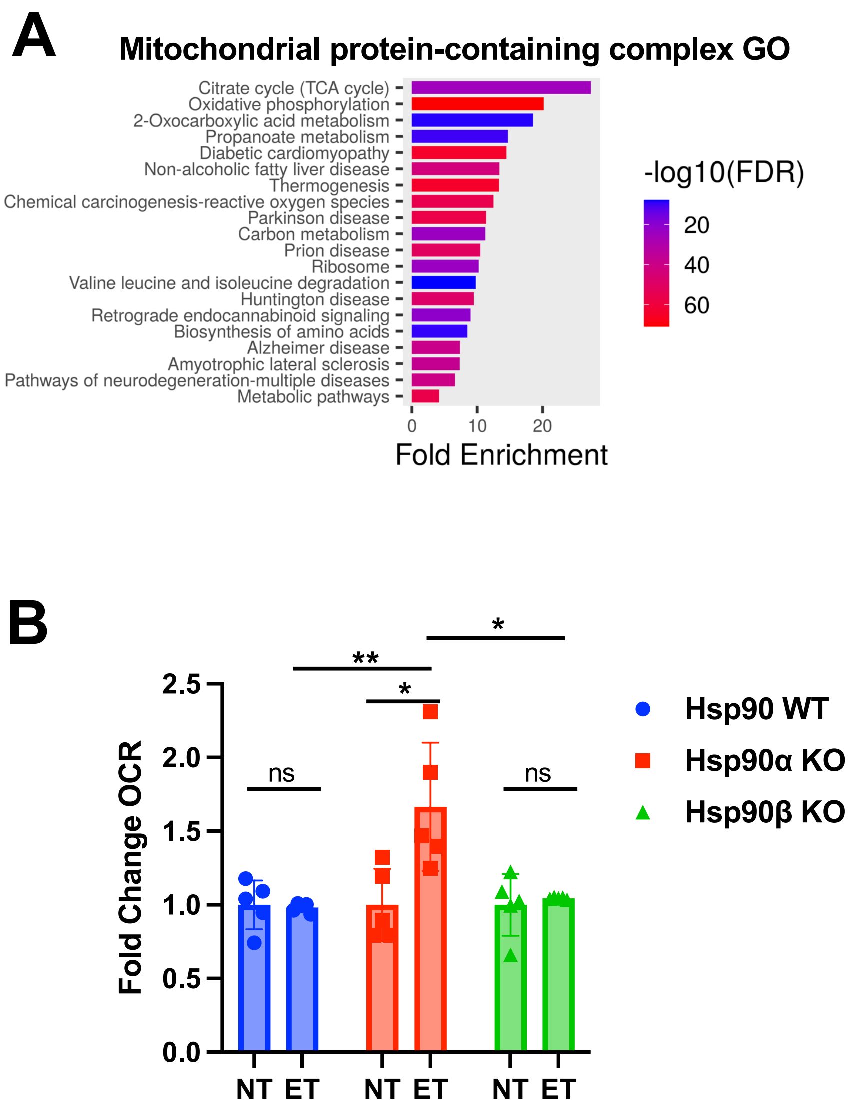

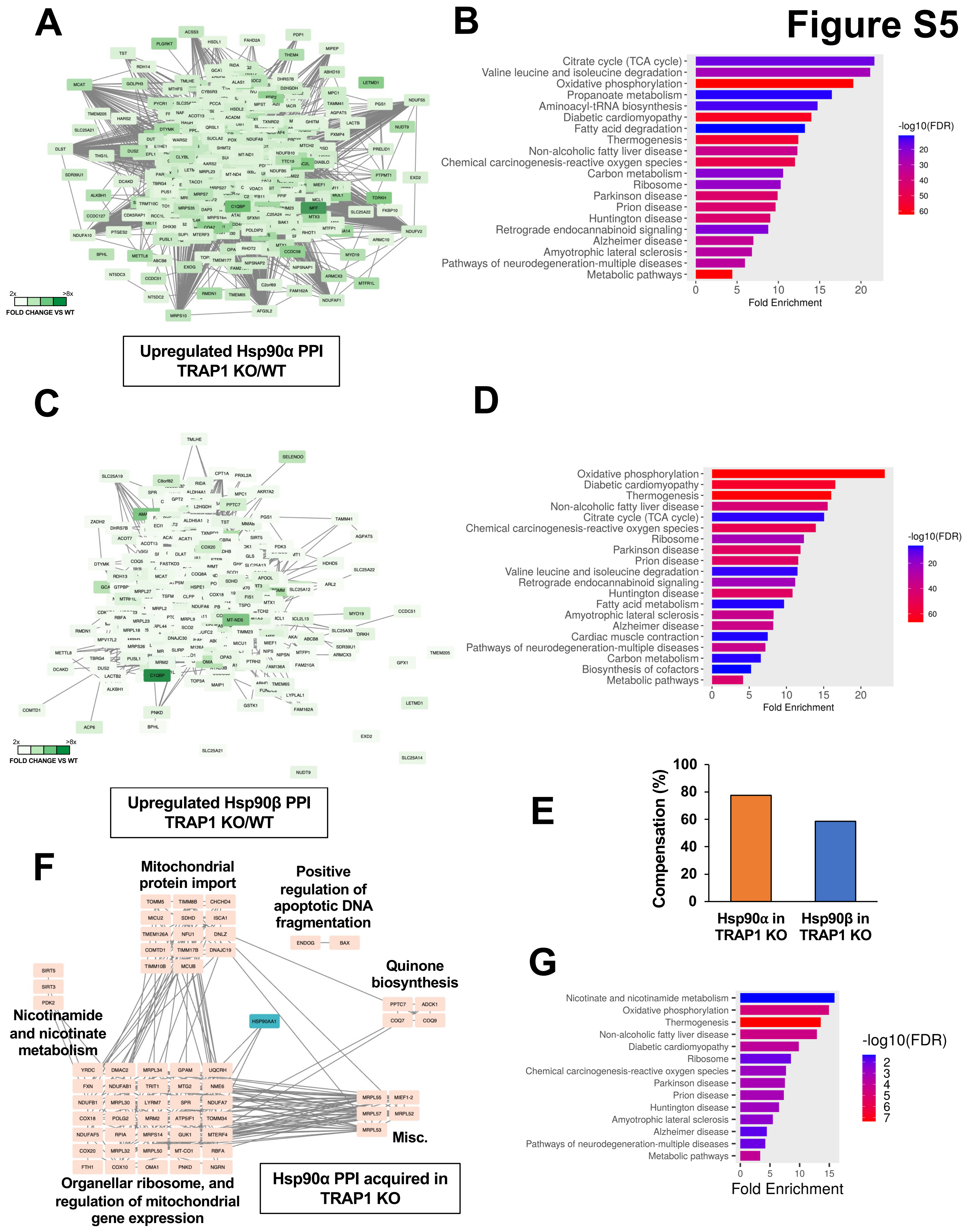

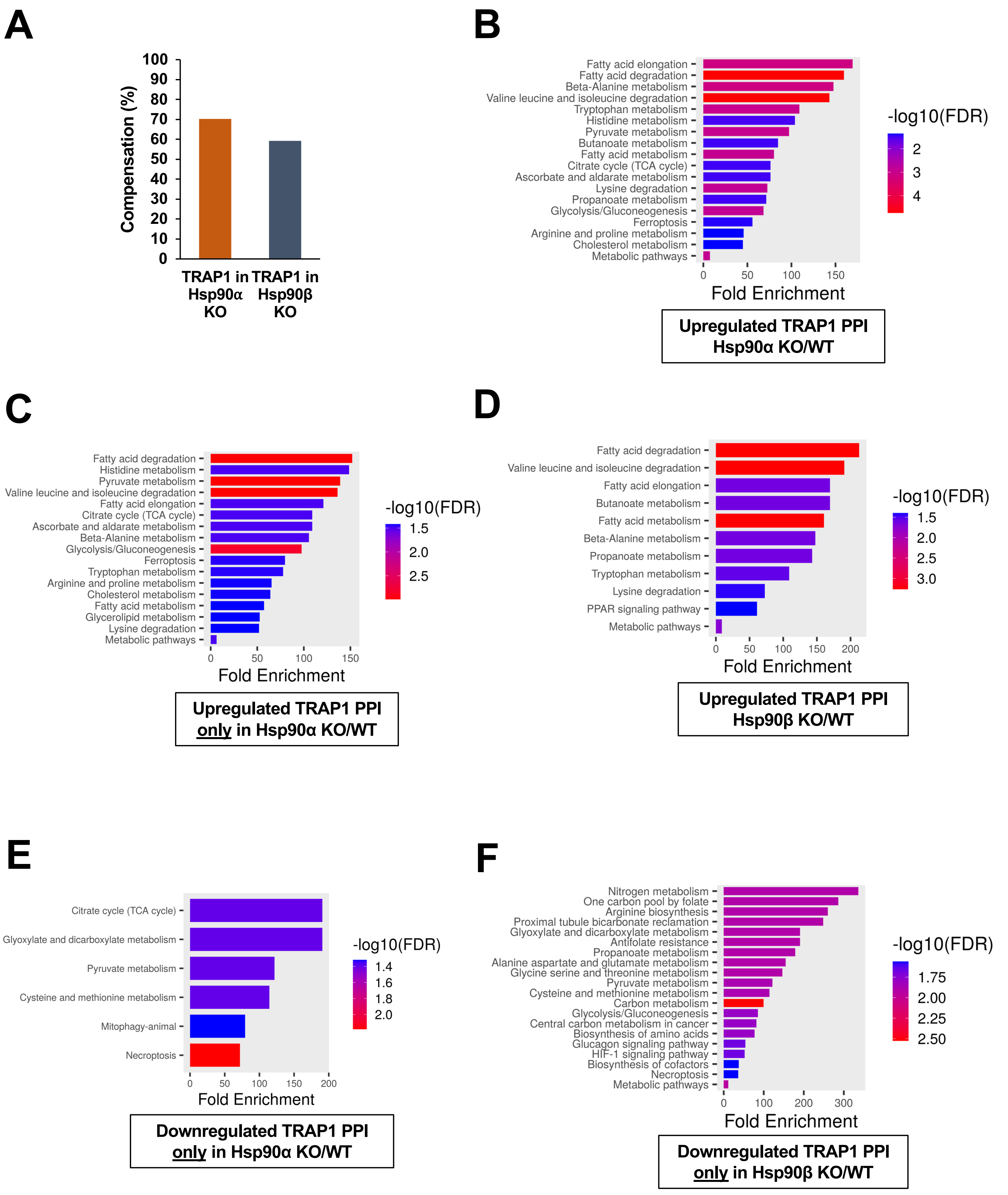

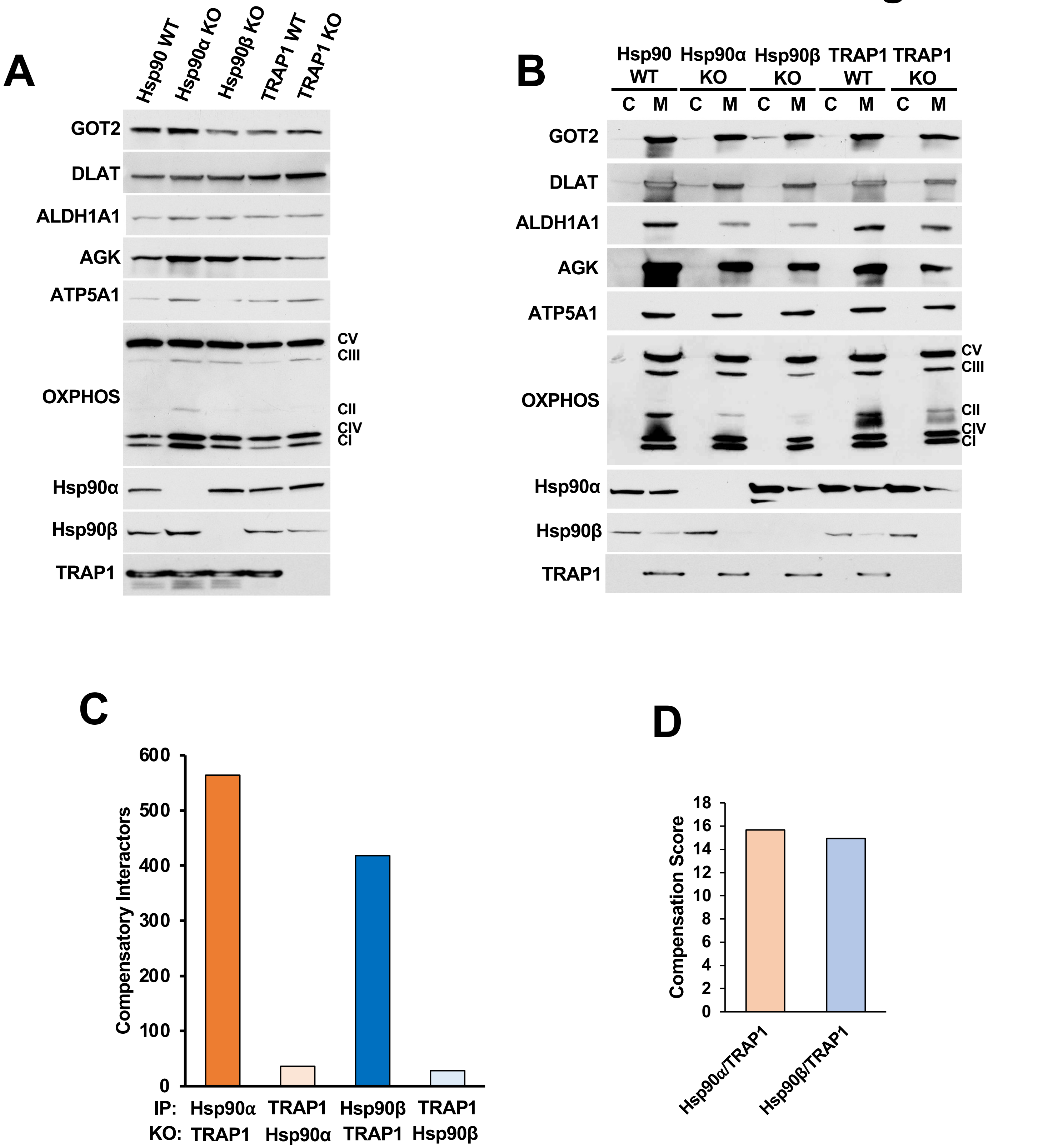

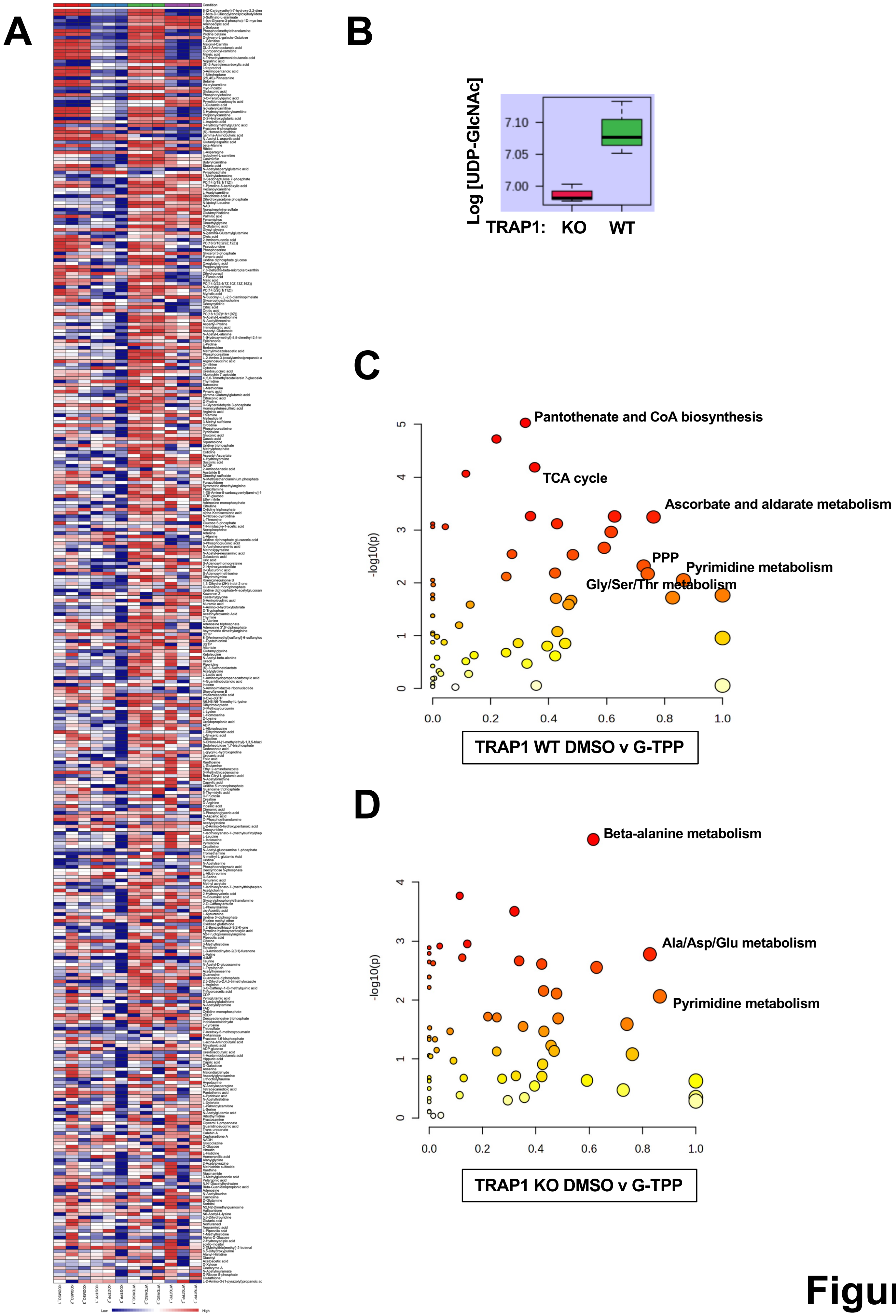

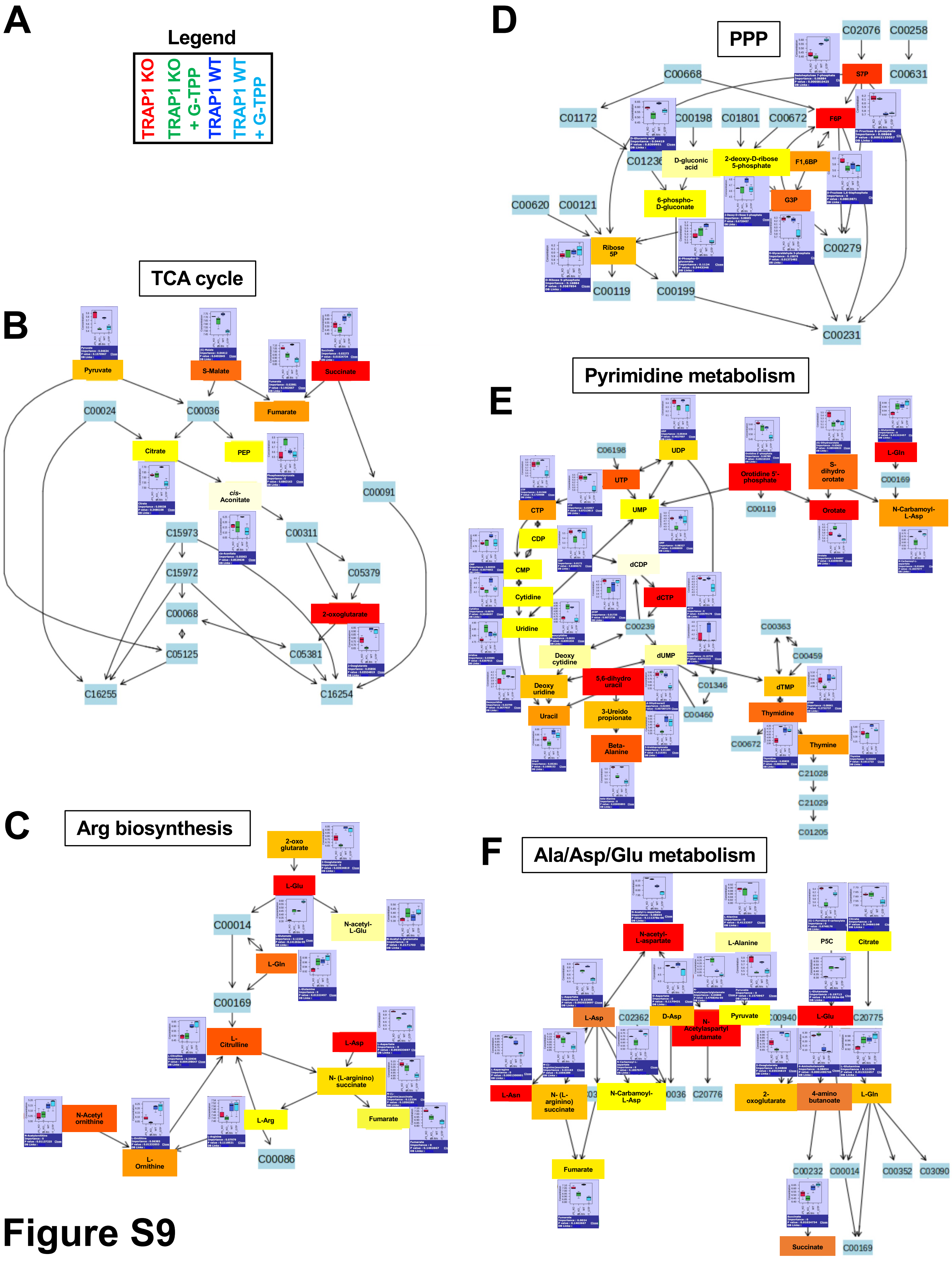
